# Alternating selection for dispersal and multicellularity favors regulated life cycles

**DOI:** 10.1101/2022.10.14.512267

**Authors:** Julien Barrere, Piyush Nanda, Andrew W. Murray

## Abstract

The evolution of complex multicellularity opened paths to increased morphological diversity and organizational novelty. This transition involved three processes: cells remained attached to one another to form groups, cells within these groups differentiated to perform different tasks, and the groups evolved new reproductive strategies^1–5^. Recent experiments identified selective pressures and mutations that can drive the emergence of simple multicellularity and cell differentiation^6–11^ but the evolution of life cycles, in particular, how simple multicellular forms reproduce has been understudied. The selective pressure and mechanisms that produced a regular alternation between single cells and multicellular collectives are still unclear12. To probe the factors regulating simple multicellular life cycles, we examined a collection of wild isolates of the budding yeast, *S. cerevisiae*^12,13^. We found that all these strains can exist as multicellular clusters, a phenotype that is controlled by the mating type locus and strongly influenced by the nutritional environment. Inspired by this variation, we engineered inducible dispersal in a multicellular laboratory strain and demonstrated that a regulated life cycle has an advantage over constitutively single-celled or constitutively multicellular life cycles when the environment alternates between favoring intercellular cooperation (a low sucrose concentration) and dispersal (a patchy environment generated by emulsion). Our results suggest that the separation of mother and daughter cells is under selection in wild isolates and is regulated by their genetic composition and the environments they encounter and that alternating patterns of resource availability may have played a role in the evolution of life cycles.

**Visual abstract:** 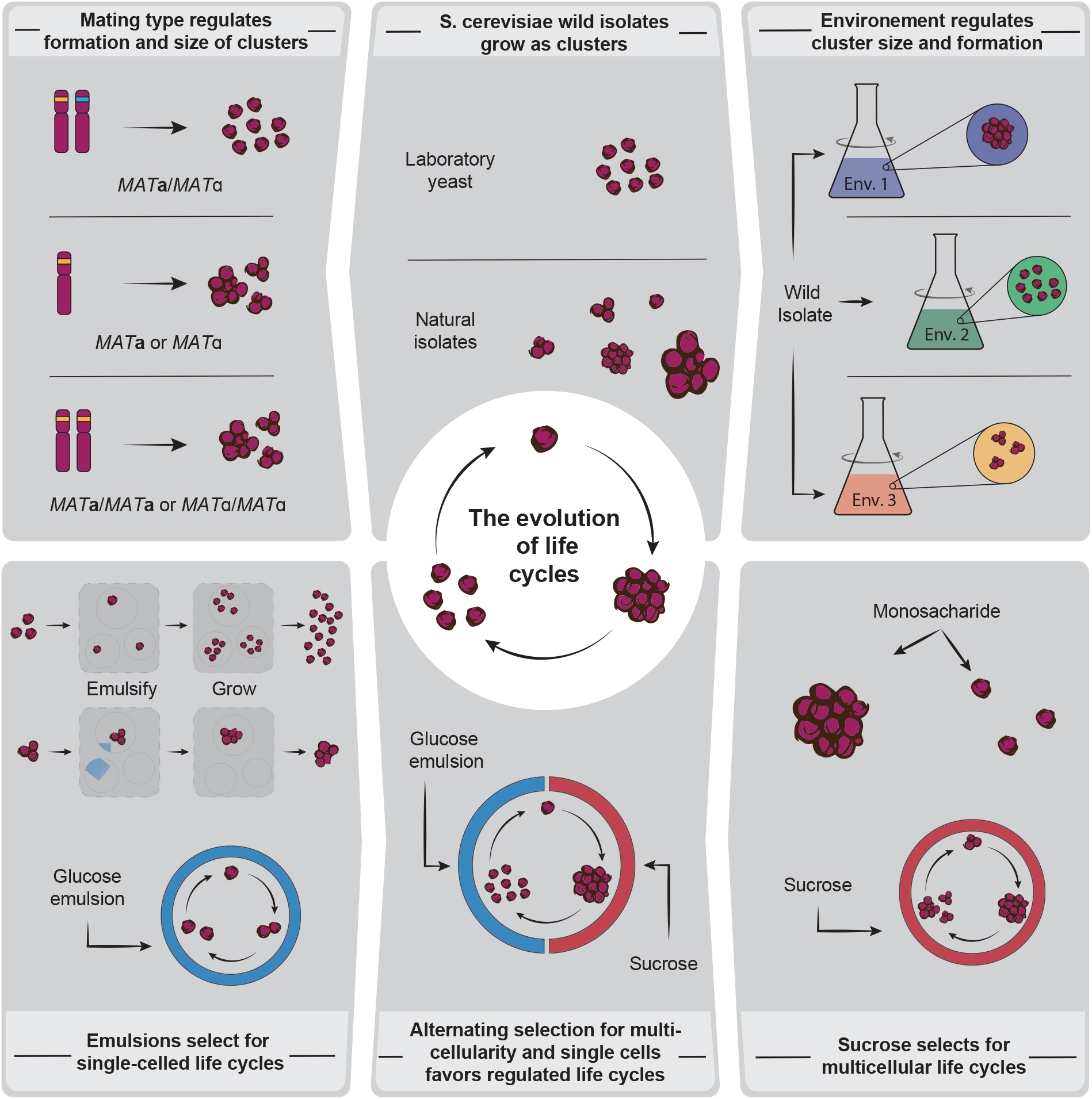

## Results and Discussion

During the evolution of multicellularity, single cells started forming clusters of multiple cells. At this early stage, multicellular life consisted of small groups of undifferentiated cells^1,2,14^. These simple groups formed either clonally or by aggregation and reproduced by randomly breaking into two or more smaller clusters, creating a minimal life cycle^1–3,5,15,16^. Here we restrict the definition of life cycle, in the context of multicellularity, to phenotypic alternation between single-celled and multicellular states^17,18^. While early life cycles might have been rudimentary, more complex life cycles have evolved resulting in a plethora of reproductive strategies with different group formation mechanisms, group features (size, shape, etc.), and propagation modes (propagule size and the signal, mechanism, and timing of offspring release)^18^. Here we explore life cycle variation within a species and the factors influencing these life cycles. We then ask which conditions could select for the evolution of regulated life cycles where multicellular growth is followed by a phase of dispersal into single cells.

Theory^17,19–25^ and empirical studies^26–28^ on the evolution of life cycles support the idea that alternating between selection for multicellularity and selection for dispersal could result in the evolution of regulated multicellular life cycles. Patchily distributed resources have been proposed as a factor that could select for dispersal and thus favor the production of single-cell propagules^29^: at the same total number of cells, single cells can colonize more resource patches than multicellular clusters. However, these ideas have not been experimentally tested, and few studies have addressed the factors influencing multicellularity within natural isolates of a single species.

Here, we characterized wild isolates of the budding yeast, *S. cerevisiae*, and showed that they form clonal multicellular clusters as part of their life cycles. These experiments also revealed that both the mating type locus and the identity of nutrients regulate the presence and size of multicellular clusters. Inspired by the environmental variation, we used a laboratory strain, W303, to engineer and compete three life cycles: constitutively single-celled, constitutively multicellular, and a regulated alternation between single cells and multicellular clusters. The single-celled life cycle was most fit in a patchy environment, the multicellular life cycle was fittest when the extracellular hydrolysis of sucrose selected for intercellular cooperation, and the regulated life cycles was fittest when conditions fluctuated between these two environments.

### The mating type locus regulates *S. cerevisiae’s* multicellularity

Laboratory strains of *S. cerevisiae* were selected to be unicellular when they were domesticated, a feature that makes them useful for studying evolutionary forces that can select for the evolution of multicellularity^6,7^. We examined wild isolates to determine the effect of genetics and the environment on simple forms of multicellularity. Here we focus on clonal multicellularity (groups formed by persistence of the shared cell wall that links mother and the daughter cells), given its widespread presence in complex multicellular organisms. To characterize life cycles, we focused on 1) quantifying the size and size distribution of multicellular clusters, and 2) identifying factors that influence cluster size.

*S. cerevisiae* can grow as clonal, undifferentiated multicellular clusters^7,30,31^. This simple multicellular phenotype has been observed in both genetically engineered and laboratory evolved strains^6,7,31^. Clusters have also been described in the wild isolates RM11-a and YL1C^32,33^, prompting us to ask if this phenotype is also present in other *S. cerevisiae* isolates. We examined 22 phylogenetically diverse wild strains from the SGRP-2 collection, sampled from various environments (wine, sake, soil, baking, etc.)^12,13^. As a control, we included *S*288c, the unicellular lab strain whose sequence is the *S. cerevisiae* reference genome.

We began by validating forward scatter, in flow cytometry, as a measure of cross sectional area and as a tool to distinguish single cells from clusters. We verified that the forward scatter of polystyrene beads is linearly correlated with their measured cross sectional area (Figure S1A) and compared forward scatter measurements of single cells to strains known to form clusters. Cultures of these clustering strains are known to include clusters of different sizes as well as single cells^7^. In addition, the area of a cluster is influenced both by the number and the size of cells it contains. To detect how large clusters can get, we developed a “clustering score” that is the ratio of the mean forward scatter of the 10% largest objects in the population to the mean forward scatter of the 10% smallest objects (Figure 1A). By focusing on the 10% largest objects, this score identifies if clusters are formed in a population. Dividing by the smallest 10% of the objects, which in most strains are almost all single cells, corrects for the differences in cell size between environments and strains (Table S1). This clustering score is therefore a relative value to compare cluster formation in different conditions and genetic backgrounds. We validated this score using diploid and haploid single cells as well as haploid clusters: single cells of different sizes show similar clustering scores, while clusters have a higher value (Figure 1A). To ensure that we were only detecting clonal multicellularity, we prevented aggregative multicellularity (often referred to as flocculation) by using ethylenediaminetetraacetic acid (EDTA) to chelate divalent cations before measuring the clustering score^34^.

**Figure 1:**
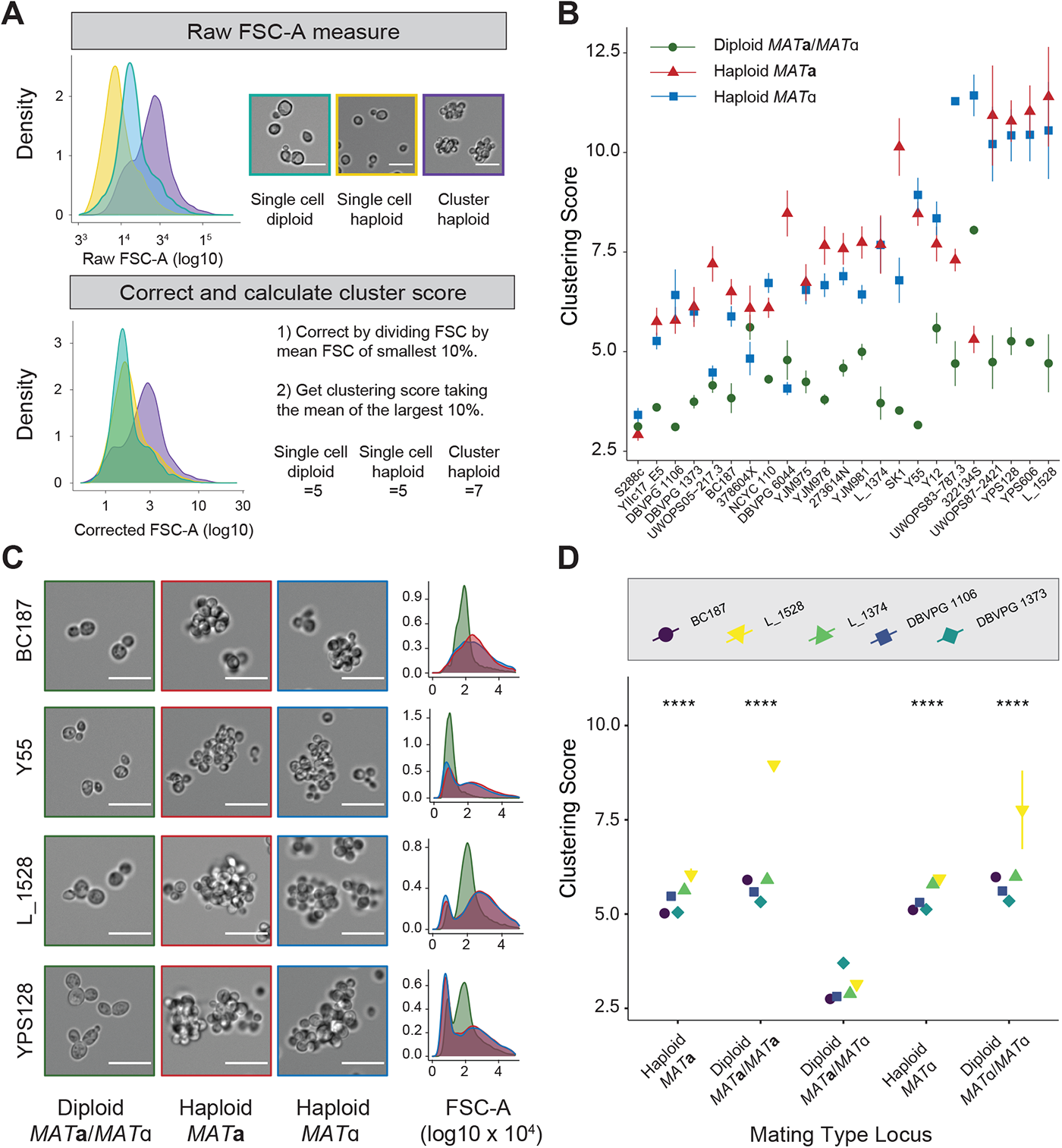
Wild S. cerevisiae isolates grow as multicellular clusters in their haploid state. A) Description of the clustering score. FSC stands for flow cytometry “Forward SCatter”. Scale bar represents 25 µm. Data for the single-celled diploid and haploid are prototrophic W303 strains. The haploid cluster is the evolved cluster Evo2 from Koschwanez et al.^7^ B) Clustering score measurements of the wild isolates grown in YPD. Haploids MAT**a** and MATα are statistically different from the diploids (p-value < 1e-13), while haploids are not different from one another (p-value = 0.3) (Wilcoxon test). Error bars represent the standard error of the mean for 3 measurements obtained from independent experiments. C) Representative DIC (Differential Interference Contrast) images and forward scatter (FSC-A) distributions of a subset of the strains (the densities on the y axis are x10^4^). Scale bar on the images represents 25 µm. D) Clustering score measurements for a subset of strains comparing haploid, diploids and diploids engineered to be homozygous at the mating type locus. Error bars represent the standard error of the mean for 3 biological replicates from the same experiment and p-values are the results of a Wilcoxon test: ****: p < 1e-4 for comparisons against the MAT**a**/MATα diploid. See also Figure S1 and Table S1.

We measured the clustering score of all 22 wild isolates and S288c both as homozygous diploids and haploids of both mating types (*MAT***a** and *MATα*) in rich, glucose-containing medium (YPD: yeast extract plus peptone (YEP) + 2% glucose). Our measurements showed that in their diploid state, the strains have low clustering scores, comparable to the values of *S*288c (Figure 1B). The haploids derived from wild isolates, however, showed a much greater clustering score, suggesting the presence of multicellular clusters (Figure 1B). Microscopy on a subset of these strains confirmed that they form multicellular clusters (Figure 1C) and reveal that natural isolates harbor a wide diversity of cluster morphology and size.

The contrasting phenotypes of haploids and diploids suggested that the mating type locus, which is heterozygous in diploids *(MAT***a***/MATα)* and present in a single copy in haploids *(MAT***a** or *MATα)*, plays a role in controlling multicellularity. We tested this hypothesis by randomly selecting 5 diploid isolates and creating derivatives homozygous at the mating type locus (*MAT***a/***MAT***a** or *MATα*/*MATα* diploids). Figure 1D shows that becoming homozygous at the mating type greatly increased the size and frequency of clusters. Our results reveal that the formation of clonal multicellular clusters in wild isolates is internally controlled at least in part by the mating type locus.

### Environment modulates multicellular life cycle in *S. cerevisiae* wild isolates

Environmental cues often drive transitions between different phases of life cycles^35^. Hence, we asked if the formation of clusters is influenced by the environment in which they grow.

We grew our strains in rich medium (YEP, yeast extract and peptone) containing five different carbon sources (glucose, sucrose, galactose, glycerol or ethanol). For each carbon source, we tested 2 different concentrations (0.2 and 2% (w/v)), and measured the clustering score in each of these environments.

Figure 2A and 2B show that the clustering score of some strains changes drastically between different environments. The effect of the carbon source was highest at a concentration of 2% (Figure 2C and Figure S2). The strongest effect was observed with the diploids in galactose: while diploids form single cells in glucose, many strains form clusters in galactose. These experiments also reveal that nutrient concentration can affect clustering. While most strains have a higher clustering core at higher concentrations of a given carbon source, a few strains show the opposite response. However, we not able to infer whether phylogenetic relationships explain variation between isolates, likely because of our small sample size (Figure S1D). Our results show the plasticity of *S. cerevisiae’s* multicellularity in response to the environment, where carbon source can act as a signal for switching between single-celled and multicellular stages of the yeast life cycle.

**Figure 2:**
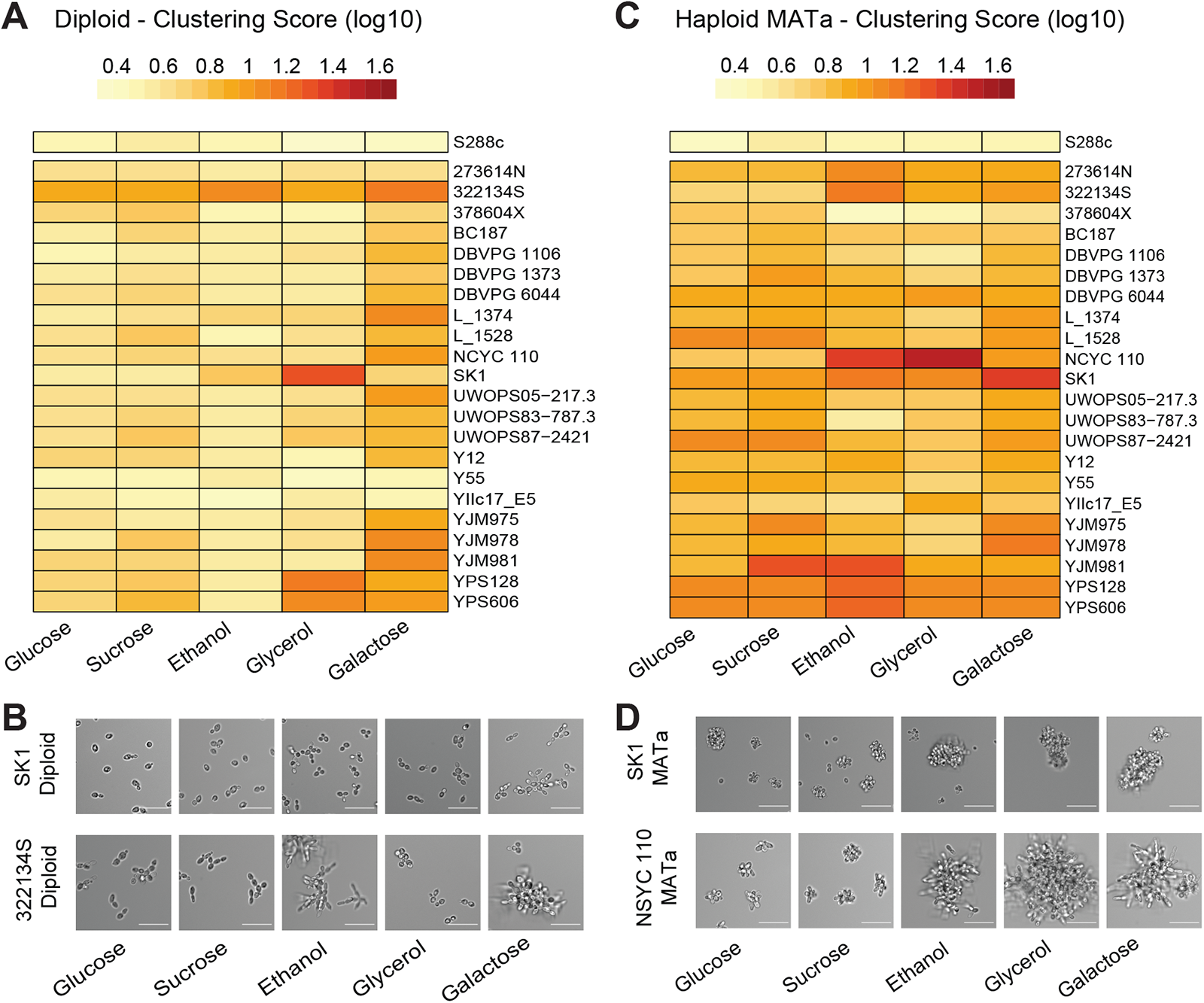
Environment modulates multicellularity in wild S. cerevisiae isolates: A) Clustering score for diploid isolates. Strains were grown in yeast extract and peptone (YEP) complemented with 5 different carbon source (concentration of 2% (w/v)). The clustering scores are the average of 3 measurements from independent experiments. B) Representative DIC images of selected diploid strains. Scale bar represents 50 µm C) Same as A but for MAT**a** strains. See Figure S2 for MATα heatmap. D) Same as B but for MAT**a** strains See also Figure S2.

### Fluctuation between bulk sucrose and glucose emulsion selects for regulated life cycles

Observing that the nutritional environment could influence the degree of multicellularity prompted us to ask if environmental oscillations could select for alternation between single-celled and multicellular phases of a life cycle. To address this question, we engineered the standard yeast laboratory strain, W303, previously used to study multicellularity^7,31^. Like S288C, W303 is single-celled both as a diploid and a haploid, unless it has been engineered to cluster. Using this strain, we analyzed how the environment influenced competitions between strains with three reproductive modes: 1) a constitutively single-celled life cycle; 2) a constitutively multicellular life cycle and 3) a regulated life cycle able to switch between single cells and clusters (Figure 3A).

**Figure 3:**
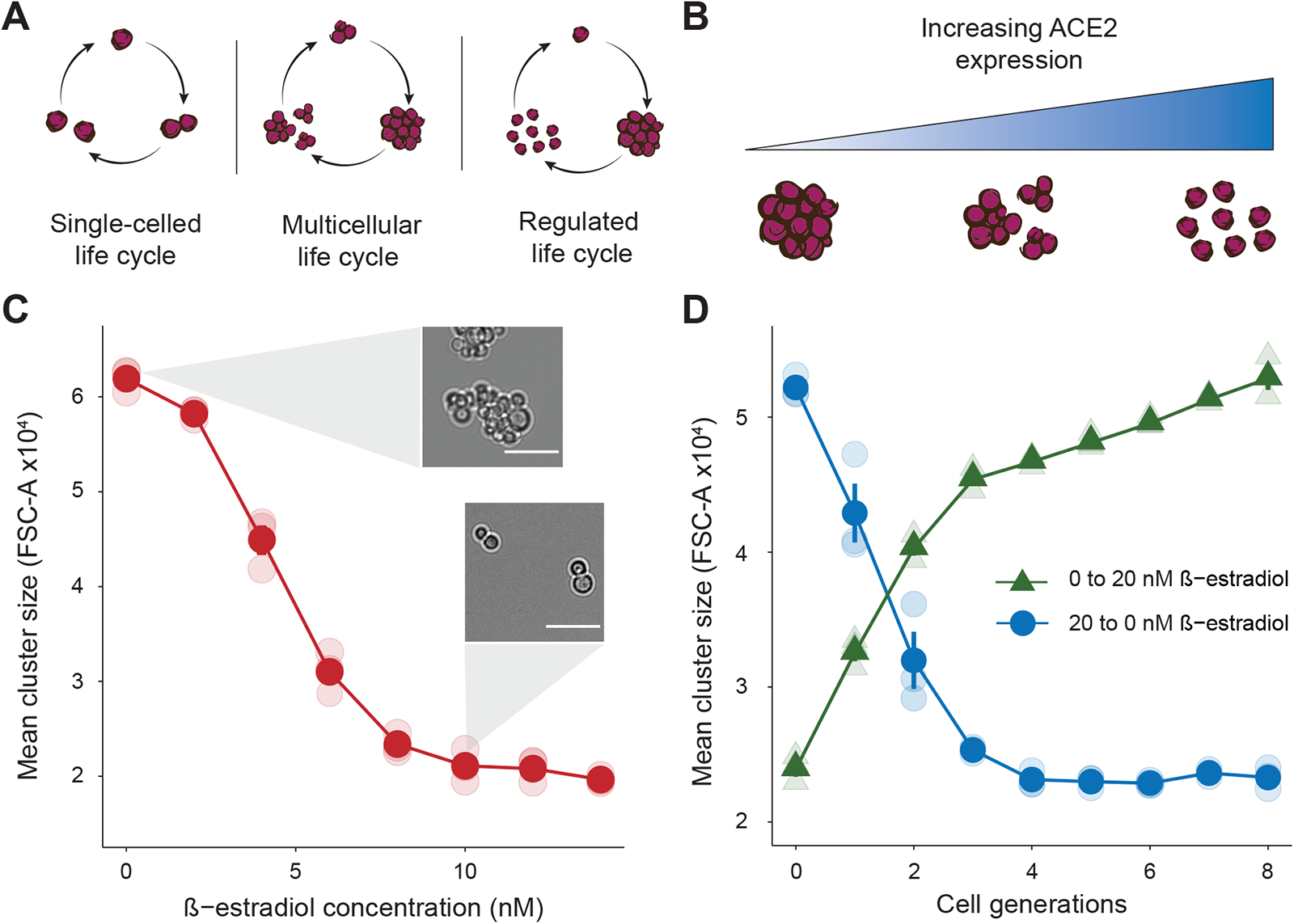
Regulated life cycles can be engineered. A) Cartoon representation of the different engineered life cycles, Single-celled life cycle: reproduction as single cells (most lab strains) . Multicellular life cycle: multicellular yeast clusters that reproduce by breaking into smaller clusters. Regulated life cycle: a strain engineered to be able to reproduce either as a single cell, or a multicellular cluster. B) To create a regulated life cycle, we placed the ACE2 gene under a promoter regulated by β-estradiol. In the absence of β-estradiol, ACE2 is not transcribed, and the strain reproduces as clusters. Increasing the β-estradiol concentration produces a single-celled life cycle. C) Mean forward scatter of the ACE2-inducible strain grown in yeast nitrogen base (YNB) with 10 mM glucose at increasing concentration of β-estradiol. Error bars (too small to be seen) represent the standard error of the mean for 3 biological replicates. Scale bar represents 50 µm. D) Kinetics of cluster formation and dissolution on the removal or addition of β-estradiol. Error bars represent the standard error of the mean for 3 biological replicates. See also Figure S3.

For the constitutively single-celled life cycle, we used a prototrophic, single-celled W303 strain. For constitutive multicellularity we deleted *ACE2*, which encodes a transcription factor that induces genes that separate daughter cells from their mothers, in the same background. Finally, to create a regulated life cycle, we placed *ACE2* under the control of a promoter that is activated by *β*-estradiol^36^ allowing the strain to grow as clusters when uninduced and as single cells in the presence of *β*-estradiol. To confirm that we could control cluster size in the *ACE2-*inducible strain, we measured cluster size at increasing *β*-estradiol concentrations, and the number of generations needed for clusters to become single cells and vice versa. Figure 3C shows that increasing *β*-estradiol decreases cluster size and Figure 3D reveals that the switches between single cells and clusters happens in a few generations.

We used the three strains to ask how different environments would affect their fitness. It has been suggested that environmental cycles (tidal, seasonal, trophic, etc.) could have driven the evolution of regulated life cycles alternating between two phenotypes (i.e. single cell and cluster)^37^. Hence, we asked if an environment cycle that alternated between favoring single cells and favoring clusters would favor a life cycle whose states were controlled by the environment.

Previous experiments^7,31^ showed that yeast clusters grow faster than single cells in low concentrations of sucrose because the extracellular hydrolysis of sucrose creates a local concentration of glucose and fructose, which can be shared between cells of a same cluster. We, therefore, used low sucrose as a selection for clusters. We tested the advantage of clusters by competing them against single cells in different concentration of sucrose. We used a mixture of glucose and fructose as a control (hexose monomer concentration ranging from 10 to 40 mM). We also confirmed the effect of dilution at each transfer during the competition experiment as group benefit is expected to decrease at high population density, when glucose and fructose can be accessed by non-kin cells^7,31^. As suggested by earlier work^31^, these experiments showed that clusters, grown on sucrose, have a fitness advantage over single cells and that this advantage is stronger as the sucrose concentration decreases (Figure S4A). We also confirmed that the dilution regime (strength of dilution between transfers of the competition experiment) greatly influences the fitness. Dilutions lower than 10,000-fold do not result in any fitness differences between single cells and clusters (Figure S4B).

These and previous results^7,31^ suggest that increased cluster size should correlate with an increased fitness on low sucrose concentrations. To directly test this hypothesis we measured the fitness of the inducibly multicellular strain expressing different levels of Ace2, allowing us to directly measure fitness as a function of size (Figure 4A). Figure 4A shows a positive relationship between size and fitness in low sucrose, confirming that size directly correlates with fitness advantage in low sucrose. In the control, where a mixture of glucose and fructose replaced sucrose, there was no difference in fitness as a function of size as the clusters are too small to limit the diffusion of nutrients to the center of the cluster^38^.

**Figure 4:**
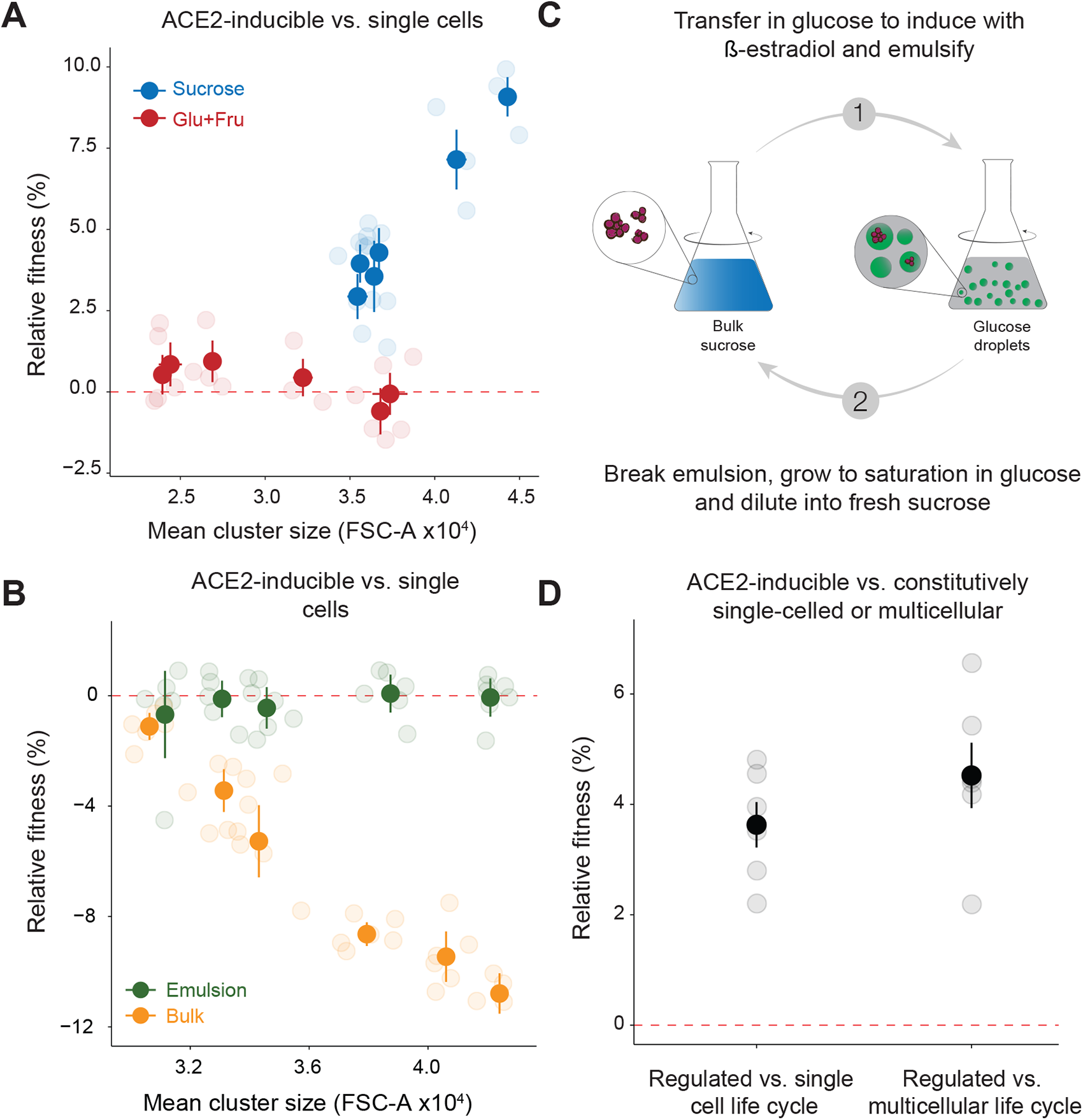
Resource availability controls the fitness differences between different life cycles. A) Competition experiment between the ACE2-inducible strain and a constitutively single-celled reference strain in YNB with 10 mM sucrose (blue) or YNB with 10 mM glucose and 10 mM fructose (red) as a control. Each point shows the fitness of the inducible strain relative to the constitutively single-celled strain at a given ACE2 induction level. FSC-A stands for Forward SCatter Area and correlates with cluster size. Competitions performed for 3 cycles of growth and dilution. Error bars represent the standard error of the mean for 3 biological replicates. B) Competition experiment between the ACE2-inducible strain and a constitutively single-celled reference strain in YNB with 100 mM glucose in emulsion (green) or in bulk as a control (grey). Each point shows the fitness of the inducible strain relative to the constitutively single-celled strain at a given ACE2 induction level. Error bars represent the standard error of the mean for 3 biological replicates. Cells were seeded at density that corresponded to a mean of less than 1 cell per drop of the emulsion. Competitions performed for 3 cycles of growth and dilution. C) Protocol for the fluctuating selection regime. Cells are first grown for a cycle in YNB with 10 mM sucrose before being transferred to YNB with 100 mM glucose and induced with β-estradiol for 24 hours. After induction, cells are put in an emulsion of YNB with 100 mM glucose for 48h. The emulsion is then broken, and cells are allowed to recover in 100 mM glucose YNB before the cycle starts again. Error bars represent the standard error of the mean for 3 biological replicates. D) Competition experiment in the fluctuating selection regime between the ACE2-inducible strain and strains that are either constitutively single-celled or constitutively form clusters. Error bars represent the standard error of the mean for 3 biological replicates. Competitions were performed for 2 full selection cycles. See also Figure S4.

To favor single cells, we created a patchy environment, in which resources were concentrated in small, separated patches by emulsifying glucose-containing medium in an inert oil. When populations are seeded in this environment, at a density of less than one cell per glucose-containing droplet, single cells should outcompete clusters thanks to their dispersal advantage: a cluster of 10 cells can only colonize 1 patch, while 10 individual cells can colonize 10 patches. Because the yield in a patch is set by the amount of the limiting resource in the patch, not the number of cells that invade it, more colonized patches result in a greater final population size.

We used an emulsion, whose droplets contained glucose and fructose, as a patchy environment. The starting cell density was adjusted so that on average a droplet does not contain more than one cell, which means that clusters will seed a much smaller fraction of the patches than single cells. The cultures were grown to saturation for 48 h in emulsion before breaking the emulsion, diluting the culture and starting a new cycle of emulsion growth. We confirmed the stability of the emulsion by measuring the size of droplets just after emulsification and after 48h (Figure S4D).

In addition, we confirmed that the emulsification protocol itself did not bias the frequency of single cells and clusters: we broke the emulsion immediately after its creation and measured the frequency of single cells and clusters (Figure S4C). Figure 4B shows that as the size of the clusters increases their fitness relative to single cells decreases. As a control, we performed the same experiments in a homogeneous and well-stirred environment and did not observe any fitness difference as a function of size.

With two environments, one favoring single cells and one favoring clusters, we could finally test whether fluctuation between them would favor regulated life cycles. We therefore competed our inducible strain against single cells or clusters in an alternating selection regime. Cells were grown for one cycle of growth in low sucrose followed by a cycle in the glucose emulsion before being transferred back in sucrose for the cycle to start again. To simulate a regulated life cycle, we induced our *ACE2* inducible strain to break the clusters into single cells before they entered the emulsion and allowed them to grow as clusters for the remainder of the competition experiment, reflecting an ideal scenario in which alternating environments regulate the stages of a life cycle (Figure 4C). Figure 4D shows that this fluctuating environment caused regulated life cycles to outcompete both single-celled and multicellular life cycles, confirming our hypothesis. Directly competing the 3 life cycles together, also confirms that the regulated life cycle outcompetes the two other life cycles (S4 E and F).

Taken together, these fitness assays show that the outcome of competition between clusters and single cells is determined by the environment: in the homogenous, low sucrose environment, clusters are fitter, but in a patchy resource environment, it is single cells that are fitter. Most provocatively, our results suggest that alternating selection for dispersal and multicellularity favors regulated life cycles.

Regulated, multicellular life cycles have evolved, independently, in many branches of the tree of life, including the fungi^1,4,18^. Many of these cycles pass through single-celled intermediates. We examined wild isolates of the budding yeast, *Saccharomyces cerevisiae*, and found that they form clonal multicellular clusters whose size is regulated by the mating type locus and their environment. Observations of larger clustering score in higher concentrations of carbon sources suggest that growth rate may contribute to cluster size regulation. The transient presence of simple multicellularity and its genetic and environmental regulation suggested that *S. cerevisiae* may have a more complex life cycle than previously thought. These results prompted us to ask if a regulated multicellularity confers an advantage over being constitutively single-celled or constitutively multicellular. We engineered a lab strain to exist in three states: constitutively single-celled, constitutively multicellular and capable of an environmentally regulated life cycle. Competing these strains with each other reveal that the single-celled life cycle is favored in a patchy resource environment, multicellular clusters are favored in an environment requiring metabolic cooperation, and that an alternation between these two environments favors a regulated life cycle.

Clonal multicellularity in yeast is due to the failure to degrade the primary septum, the specialized portion of the cell wall that holds mother and daughter cells together after cytokinesis has separated their cytoplasms. The increased cluster size in haploids strains suggests that the expression of haploid specific genes may hinder the production of enzymes that degrade the primary septum. For example Ste12, a haploid specific transcription factor, is required for the expression of *AMN1*, a gene known to inhibit cell separation and the binding of the Mat**a**1/Mat*α*2 heterodimer prevents Ste12 from accessing the *AMN1* promoter in diploid cells^33^.

Clonal multicellularity has an intrinsic advantage, resistance to cheating, over the agglomerative multicellularity exhibited by *Dictyostelium discoideum* or *Myxococcus xanthus*^1,2,39–42^. However, the group formation mechanisms constrain clonal multicellularity’s ability to rapidly switch from single cell to multicellularity. Under the fluctuating regime selection described here, we would predict that aggregative multicellularity carries an advantage as the switch from single cell to multicellularity can happen in less than a generation.

Most natural isolates of *S. cerevisiae* are diploid and these diploids are primarily formed by mating between sister spores produced from the same meiosis. The budding pattern of haploid cells has been proposed to favor the mating that occurs when a germinated spore lacks an appropriate partner and uses mating type switching to produce descendants that have opposite mating types^43,44^. By forming multicellular clusters, these clonally related cells would still be able to mate, even in environments where fluid flow and other physical forces would otherwise separate the cells.

In a single environment, the clonal clusters produced by experimental evolution or found in natural isolates usually break into smaller clusters rather than giving rise to populations of single cells. This mode of reproduction is favored in environments where metabolic cooperation or direct selection for cluster size gives clusters higher fitness than single cells. But if the environment alternates between favoring clusters and single cells, life cycles that regulate propagule size in response to the environmental fluctuation should be favored. We created this fluctuation by alternating between an environment that selected for metabolic cooperation and one that favored dispersal in a patchy environment. Our engineered strains add evidence that experimentally constructed, alternating environments can select for a regulated life cycle^27,45^. Outside the lab, natural oscillations, such as tidal, seasonal, or ecological fluctuations could have selected for the evolution of regulated life cycle. Examining wild isolates in their natural habitats and correlating their phenotype with fluctuations in their environments, over time and space, may reveal that *S. cerevisiae* is capable of more sophisticated and ecologically adaptive life cycles than have been discovered so far.

## Acknowledgements

The authors thank members of the Murray lab, especially Thomas LaBar and Tanush Jagdish for discussions, John Koschwanez for help with developing the project and Michael Desai, Michael Laub, Silvia De Monte, and William Ratcliff for their feedback on the manuscript. J.B, P.N and AWM are supported by the grant NIH/NIGMS R01 GM043987. A.W.M. is also supported by the NSF-Simons Center for the Mathematical and Statistical Analysis of Biology (NSF #1764269, Simons #594596).

## Author contributions

Conceptualization, J.B., and A.W.M.; Methodology, J.B., and A.W.M.; Investigation, J.B. and P.N.; Writing – Original Draft, J.B.; Writing – Review & Editing, J.B., P.N., and A.W.M.; Funding Acquisition, A.W.M; Supervision, A.W.M.

## STAR Methods

### RESOURCE AVAILABILITY

#### Lead Contact

Further information and requests for resources and reagents should be directed to and will be fulfilled by the Lead Contact, Andrew Murray.

#### Materials availability

Yeast strains created in this study are available upon request from the Lead Contact.

#### Data and code availability

All protocols, data and analysis codes in this study are available at https://doi.org/10.7910/DVN/EHJYXR.

### EXPERIMENTAL MODEL AND SUBJECT DETAILS

#### Yeast strains and media

All experiments with the engineered strains were conducted in minimal (no amino acids or nucleotides) synthetic media made by combining refrigerated stocks of 10X Yeast Nitrogen Base (with ammonium sulfate) from BD Difco™ with the appropriate sugar (sucrose stocks were kept at -20°C). Once made, the minimal synthetic media was stored at 4°C and protected from light.

*β*-estradiol stocks were made at a concentration of 10mM in ethanol and stored at -20°C. To obtain the final working concentration for the experiments, the *β*-estradiol stock was diluted in ultrapure water. Experiments involving the wild isolates experiments were performed in rich medium, YEP (1% Yeast-Extract, 2% Peptone) to which we added dextrose, sucrose, ethanol, galactose or glycerol at the appropriate concentration. YPD is YEP containing 2% w/v glucose.

##### Engineered strains

All engineered strains used in this study can be found in Table S2. Engineered strains were constructed by first inserting the LexA-ER-AD system at the *HIS3* locus as described by Ottoz et al.^36^. We then replaced the *ACE2* promoter with an inducible promoter containing 1 LexA binding site to create the inducible strain. *ACE2* was deleted to create the positive control: the constitutively multicellular strain. Nothing was changed for the negative control: the constitutively single-celled strain. Each of these three strains were marked with a different fluorescent protein, expressed from the *ACT1* promoter, to allow competition essays.

##### Original wild isolates

The wild isolates used are from the SGRP-2 collection created by the Liti lab^12,13^. Before performing the experiments, we confirmed the mating type of all the strains used both by mating and by PCR^46^. Inconsistencies between reported and observed genotypes of several haploid strains were corrected by combining mating type switching and sporulation. We created a new set of diploids by mating the two heterothallic haploids, allowing us to have isogenic haploids and diploids. We used all the strains from the collection that we could make available as diploids, and both *MAT***a** and *MATα* haploid*s*: 273614N, BC187, DBVPG1106, DBVPG1373, DBVPG6044, 322134S, L_1374, L_1528, NCYC110, SK1, 378604,X UWOPS05-217.3, S288c, UWOPS83-787.3, UWOPS87-2421, Y12, Y55, YIIc17_E5, YJM975, YJM978, YJM981, YPS128, and YPS606.

##### Homozygosing the MAT locus

to create diploids homozygous at the mating type locus, we transformed diploids with a plasmid carrying the *HO* gene under an inducible promoter^47^. Colonies were selected on uracil plates and allowed to lose the plasmid on YPD before being tested for mating type both by mating and PCR using methods described here^46^.

### METHOD DETAILS

Detailed protocol for each figure is available at https://doi.org/10.7910/DVN/EHJYXR.

#### Competition essays

All competition experiments were performed in YNB complemented with the appropriate carbon source. Experiments were performed in plastic culture tubes. Fluorescently labeled strains were individually pre-grown overnight in media in which the competition experiment would be performed. Strains were then mixed in 1:1 ratios and diluted in PBS before being inoculated into the appropriate medium. Cultures were passaged for three cycles of growth and frequency of each strain was measured at each transfer with a flow cytometer (Fortessa, BD Bioscience, RRID:SCR_013311, US). Because the distribution of number of cells per cluster was stable from one transfer from another we were able to estimate the relative fitness between single cells and clusters by extracting the ratio between fluorescent events (an event here can be a single cell, or a cluster of any size). The frequency of each strain was extracted with the CytoExploreR^48^ package. To quantify the relative fitness we performed a linear regression between the number of generation elapsed between each transfer and the log of the ratio between the two genotypes. The relative fitness is the slope of the regressed line. Errors bars reflect the standard error of the mean from at least 3 independent replicates. Note that the reference strain varies for different experiments and is indicated for each plot.

#### Emulsions

Emulsions were produced by mixing 150 ul of culture with 350 ul of FC-40 oil containing 2% (w/v) 008-FluoroSurfactant (Ran Biotechnologies) in centrifuge tubes (VWR 76332-074). Note that emulsion stability is influenced by the type of centrifuge tube used. The tubes were then gently tapped before being vortexed at max speed for 30 sec. The tubes were allowed to rest before being opened and sealed with parafilm for the 48h incubation time. To break the emulsion, the tubes were first spun down with a counter top centrifuge, for 30 sec, the lower oil phase was removed before the addition of 100 ul of 1H,1H,2H,1H-Perfluoro-1-octanol (Sigma Aldrich) mixed by pipetting up and down and gently turning the tube. After 10 min of rest, tubes were spun down for 30 sec on a counter top centrifuge, the lower phase was removed before gently mixing the aqueous phase and recovering the cells by pipetting.

#### Cycling experiments

For the cycling experiments (Figure 4C), the strains were first pre-grown overnight in YNB with 100 mM glucose before being mixed in a 1:1 ratio and diluted 1:100 in YNB with 100 mM glucose to start the first step of the cycle. After 24h of incubation at 30°C on a roller drum, culture were diluted 1:10,000 in sucrose for the second step and incubated for 4 days. For the third step, the cells were diluted 1:100 in 100 mM glucose and *β*-estradiol was added for induction when needed. After 24h, the culture was diluted 1:10,000 and transferred into emulsion and allowed to grow to saturation before the emulsion was broken (as described above) and the diluted 1:100 in YNB with 100 mM glucose to restart the cycle. The frequency of each strain was measure at each transfer using flow cytometry.

#### Size measurements on wild isolates

A colony of each wild isolates was inoculated in 300 ul of YPD in a 2 ml deep 96 well plate and allowed to grow overnight on a shaker at 1000 rpm. Cultures were then diluted 1:100 in the appropriate media and incubated for 48h before being transferred again 1:100 in the same media. Samples were measured in early stationary phase, after 24h or 48 depending on the media. To prevent flocculation, cultures were diluted in EDTA to a final concentration of 100 mM. Each of the 3 replicates of these measurement was perform as an independent batch.

#### Clustering Score

To allow us to compare the wild isolates size measurements we developed a clustering score allowing us to quantify the clustering level of each strain and to compare between different strains and different environment (Figure 1A). Cultures of strains that form clusters are composed of clusters of different sizes but also of single cells. We therefore divide the forward scatter measurements by the mean of the 10% smallest forward scatter values of a population (usually mostly single cells), which corrects for differences in cell size between environments and strains. We then take the mean of the 10% largest corrected forward scatter values as the clustering score. This score is used as a relative value to compare the formation of clusters by different strains in different environments. To confirm that the 10% smallest objects are mostly single cells we imaged a subset of 10 isolates and counted the number of single cells and clusters using the Fiji distribution of ImageJ^49^ (Figure S2).

#### Microscopy

All images of cells and clusters were taken in a 96-well glass-bottomed plate (Greiner bio-one, www.gbo.com) using a Nikon Ti inverted microscope (www.nikoninstruments.com) with MetaMorph software (www.metamorph.com). Contrast of images was adjusted, and images were annotated with scale bars using the Fiji distribution of ImageJ^49^. Contrast was changed for visibility only and does not impact the results. Images of the emulsion for droplet size quantification were taken in a hemocytometer (Bulldog Bio).

---

**Figure S1.**
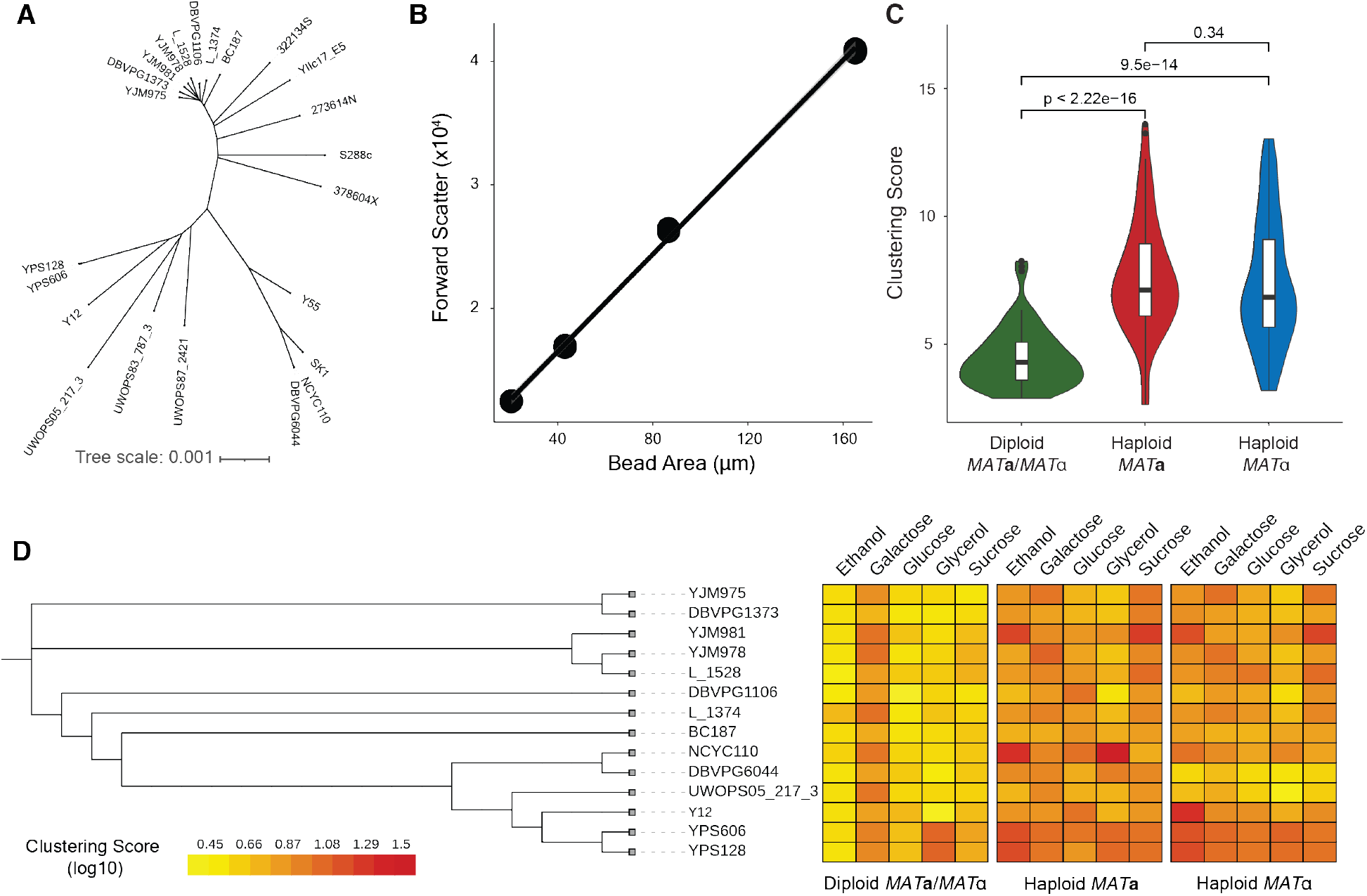
Wild isolates phylogeny and clustering score. Related to Figure 1. A) Neighbor Joining tree of the wild isolates used in this study. Trees built based on SNP differences, using data from Liti et al. 2009^13^. B) correlation between bead cross-sectional area and mean forward scatter measurements. FSC for each bead size was measured in 3 independent measurements. C) Clustering score in YPD of all 23 strains. The indicated p-values are the results of Wilcoxon test. D) Clustering score in different carbon sources arranged by phylogeny. Data restricted to the 13 strains that are not mosaics from multiple lineages.

**Figure S2.**
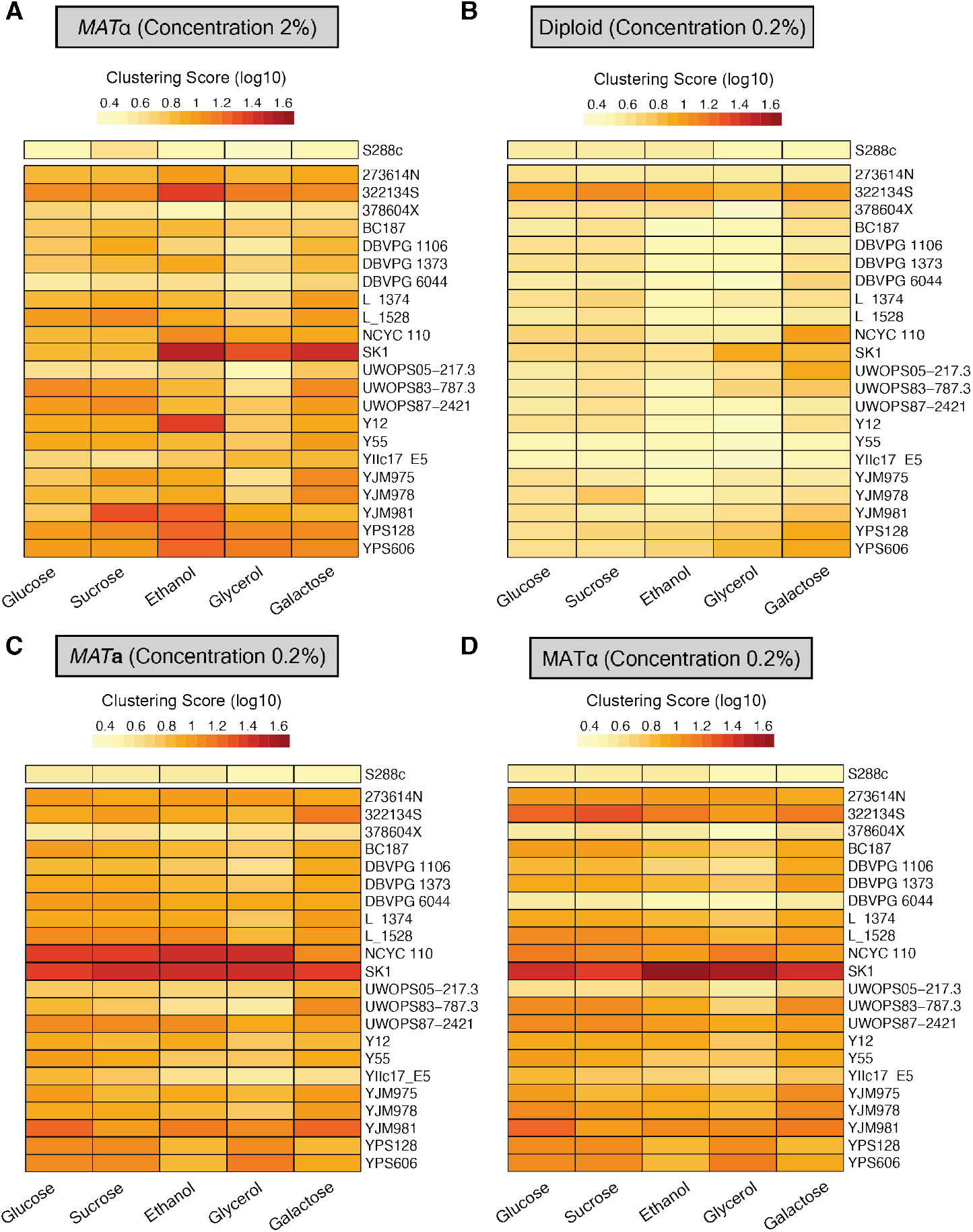
Wild isolates clustering score at different sugar concentrations. Related to Figure 2. A) Clustering score for haploid MAT α isolates. Environments are hierarchically grouped. Strains were grown in YEP complemented with 5 different carbon source (concentration of 2% (w/v)). The clustering scores are the average of 3 measurements from independent experiments. B) Same as A but for diploid strains in sugar concentration of 0.2%. C) Same as A but for MAT**a** strains in sugar concentration of 0.2%. D) Same as A but for MAT α strains in sugar concentration of 0.2%.

**Figure S3.**
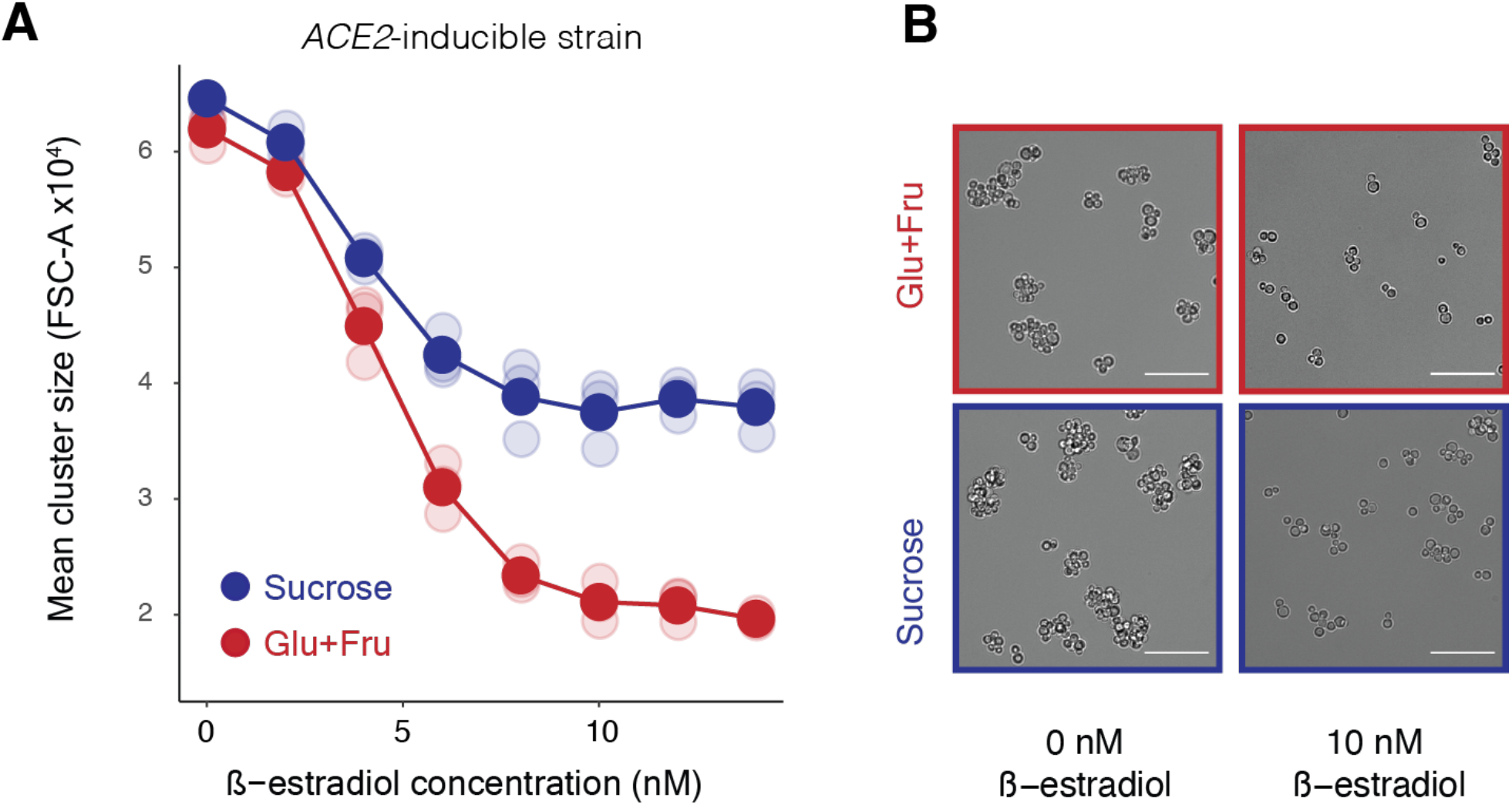
Cluster size regulation using β-estradiol. Related to Figure 3. A) Mean forward scatter of the engineered, ACE2-inducible strain in glucose + fructose or sucrose at increasing concentration of β-estradiol. Error bars, too small to be seen, represent the standard error of the mean for 3 biological replicates. B) DIC images of the ACE2-inducible strain when uninduced or induced with 10 nM of β-estradiol. Cells were grown in YNB with 10 mM sucrose or YNB with 10 mM glucose and 10 mM fructose. Scale bar represents 50 µm.

**Figure S4.**
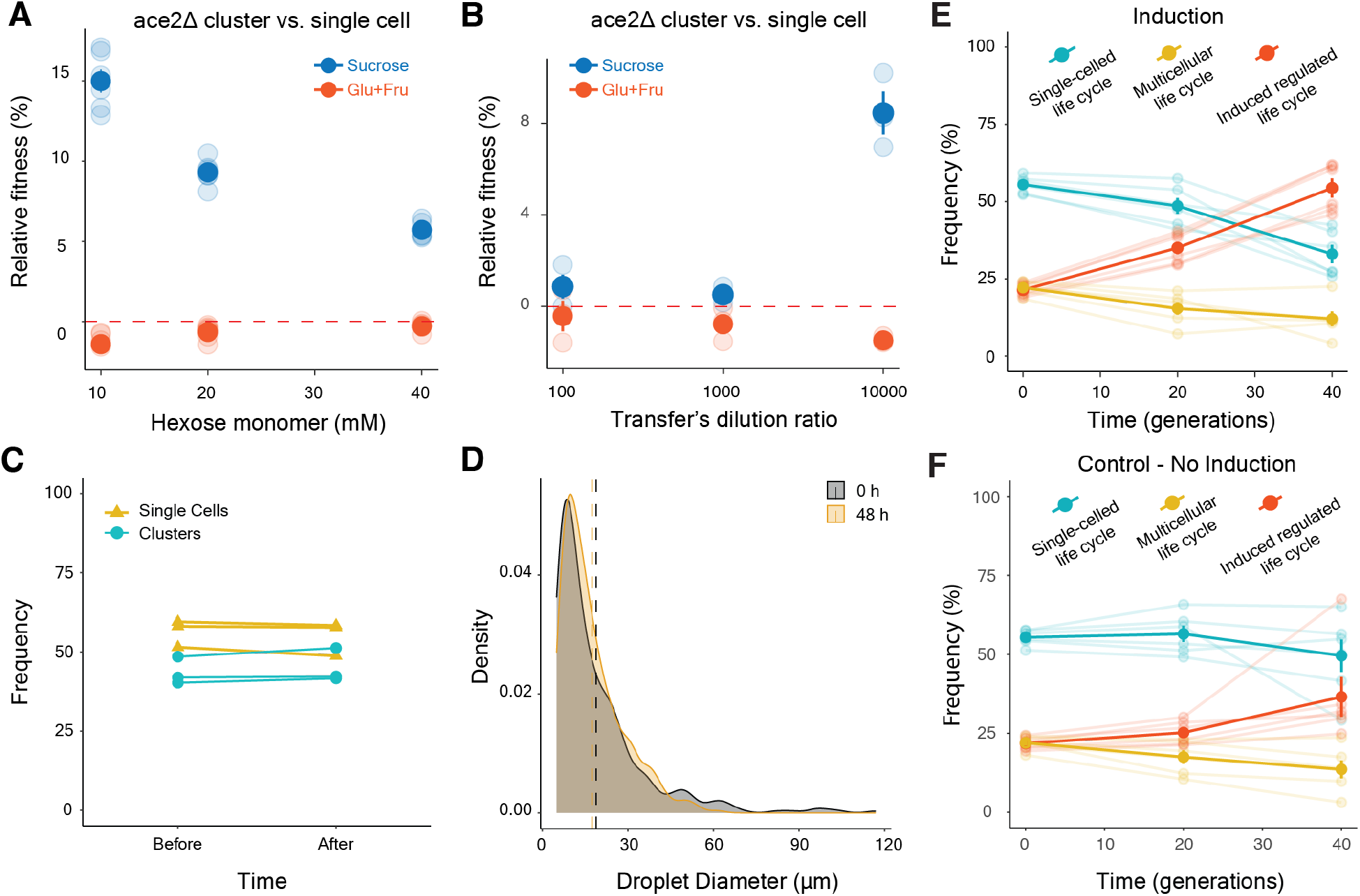
Engineered life cycles can be competed in sucrose and glucose emulsions. Related to Figure 4. A) Competition experiment between an ace2Δ cluster and a single cell in different concentrations of sucrose or glucose + fructose. These results confirm work by Koschwanez et al. (2013) and quantify the fitness advantage of clusters in sucrose. Error bars represent the standard error of the mean for 3 biological replicates. B) Similar to A but competitions were done in 10 mM sucrose and 10 mM Glucose + 10 mM Fructose with different dilution factors between transfers (100 stands for a 1:100 dilution). A large dilution factor is needed to observe fitness differences between single cells and clusters. Error bars represent the standard error of the mean for 3 biological replicates. C) Measurements of ratio of single cells to clusters before introducing them into an emulsion and after immediately breaking the emulsion, with no intervening growth, to confirm that the emulsion protocol does not create a bias in favor of clusters or single cells. D) Size distribution of emulsion droplets. The emulsion was imaged and droplet diameters were measured using Fiji. E) Frequency of the three life cycles (single-celled, multicellular and regulated induced) overtime when all were grown together in a regime alternating between bulk sucrose and glucose emulsions. The bold lines represent the means of replicates, while the light lines represent the data for individual replicates. Error bars represent the standard error of the mean for the 6 biological replicates. F) Same as E, but a control experiment where the strain designed as a regulated life cycle is not induced and therefore always grows as cluster.

**Table S1:**
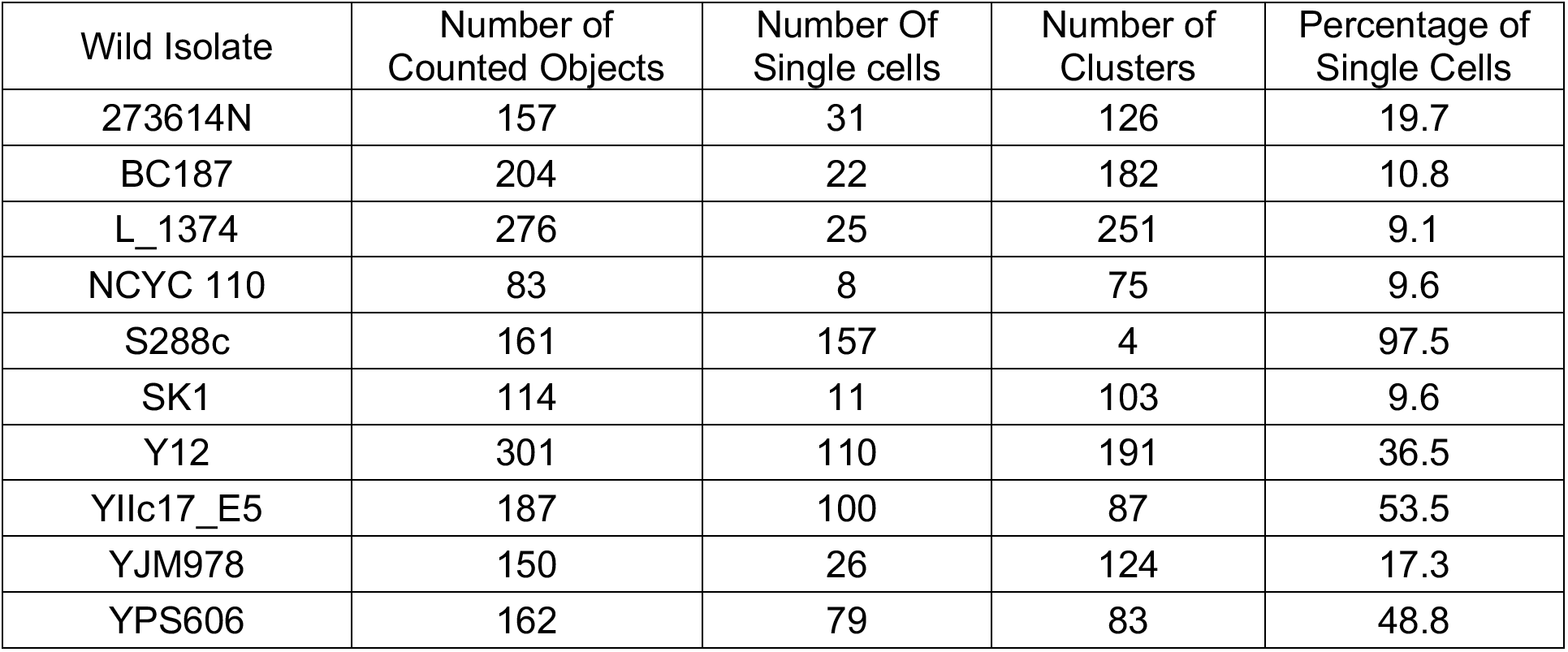
Percentage of single cells in wild isolates populations (MATa). Related to Figure 1. We randomly selected 10 wild isolates haploid MAT**a**, imaged them under the microscope and counted objects by classifying them as single cells or clusters.

**Table S2.**
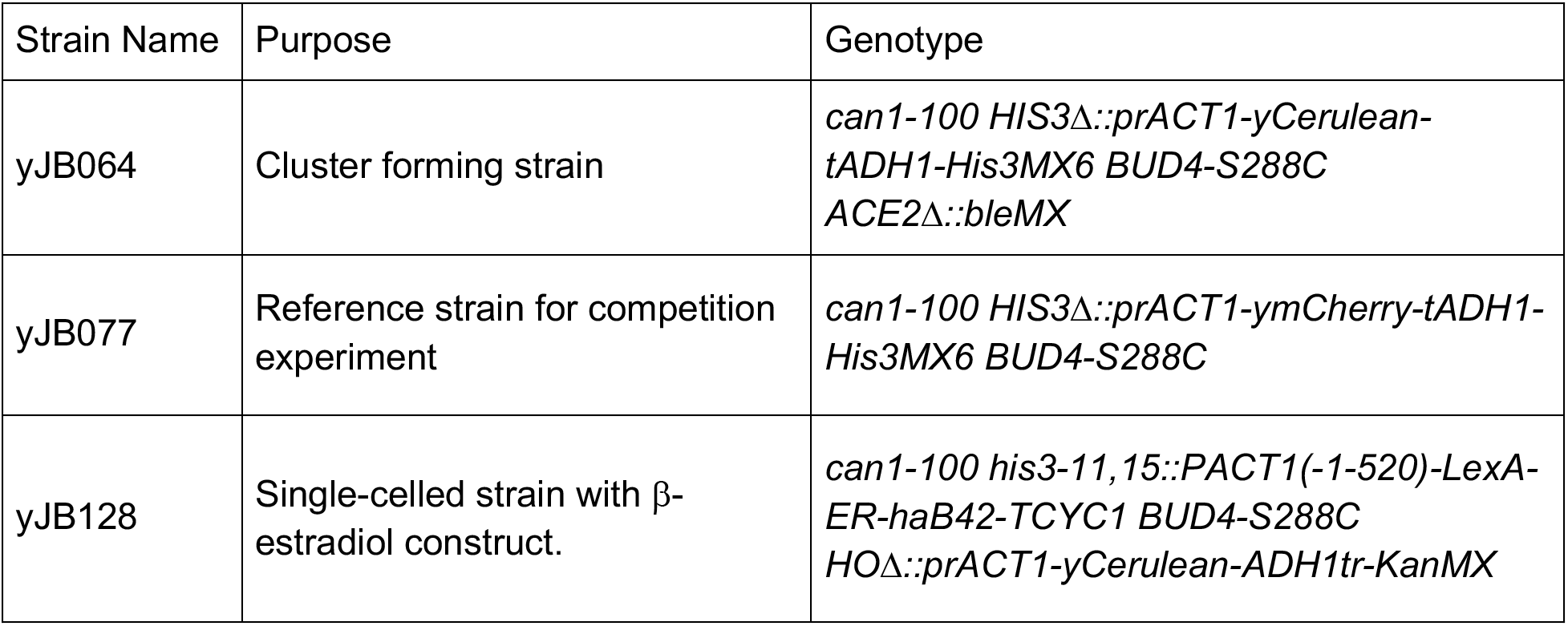
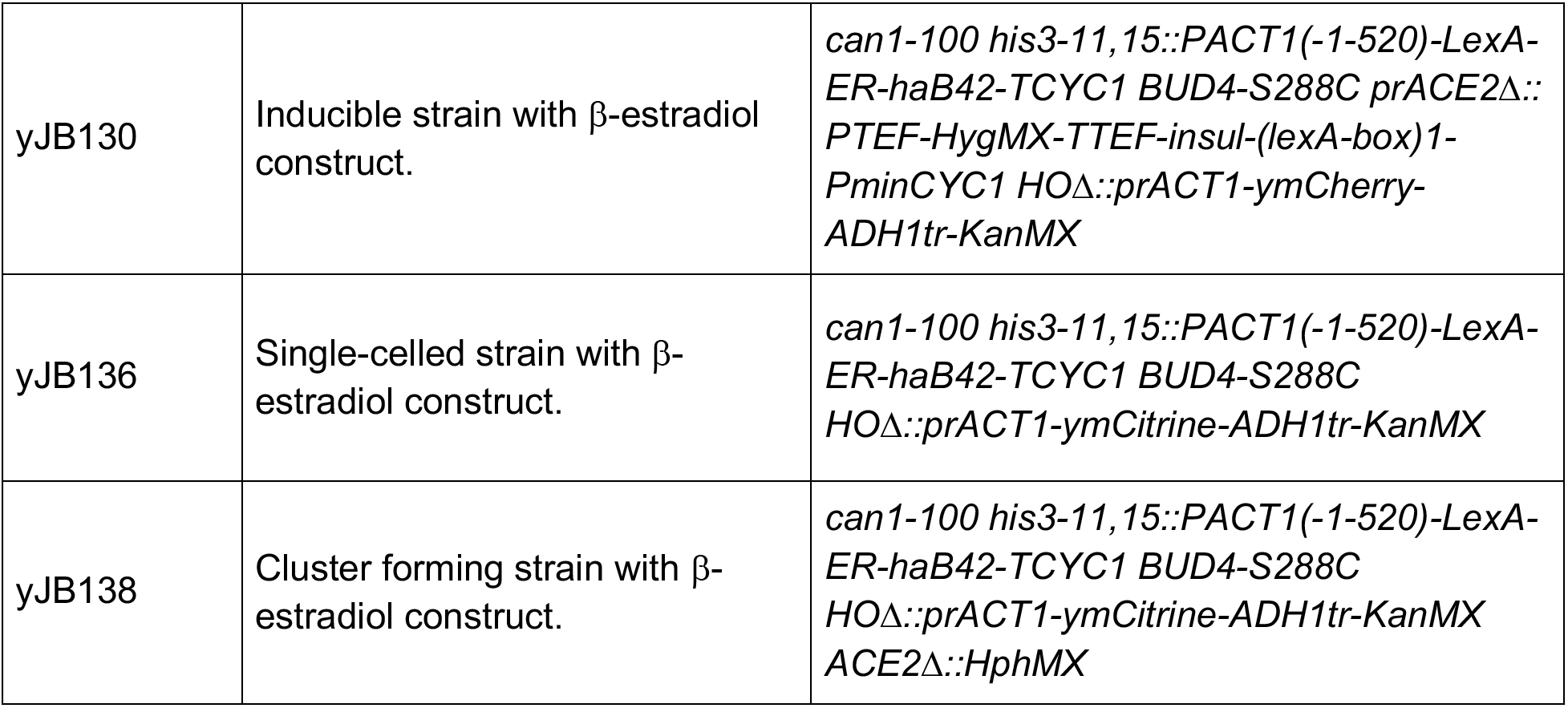
Laboratory yeast strains used in this study. Related to Figure 3 and Figure 4. All strains were derived from yJK089, a derivative of the W303 strain background

| Strain Name | Purpose | Genotype |
| --- | --- | --- |
| yJB064 | Cluster forming strain | *can1-100 HIS3Δ::prACT1-yCerulean-tADH1-His3MX6 BUD4-S288C ACE2Δ::bleMX* |
| yJB077 | Reference strain for competition experiment | *can1-100 HIS3Δ::prACT1-ymCherry-tADH1-His3MX6 BUD4-S288C* |
| yJB128 | Single-celled strain with β-estradiol construct. | *can1-100 his3-11, 15::PACT1(-1-520)-LexA-ER-haB42-TCYC1 BUD4-S288C HOΔ::prACT1-yCerulean-ADH1tr-KanMX* |
| yJB130 | Inducible strain with β-estradiol construct. | *can1-100 his3-11,15::PACT1(-1-520)-LexA-ER-haB42-TCYC1 BUD4-S288C prACE2Δ::PTEF-HygMX-TTEF-insul-(lexA-box)1-PminCYC1 HOΔ::prACT1-ymCherry-ADH1tr-KanMX* |
| yJB136 | Single-celled strain with β-estradiol construct. | *can1-100 his3-11,15::PACT1(-1-520)-LexA-ER-haB42-TCYC1 BUD4-S288C HOΔ::prACT1-ymCitrine-ADH1tr-KanMX* |
| yJB138 | Cluster forming strain with β-estradiol construct. | *can1-100 his3-11,15::PACT1(-1-520)-LexA-ER-haB42-TCYC1 BUD4-S288C HOΔ::prACT1-ymCitrine-ADH1tr-KanMX ACE2Δ::HphMX* |

