## Supplementary Figures for "Alternating selection for dispersal and multicellularity favors regulated life cycles"

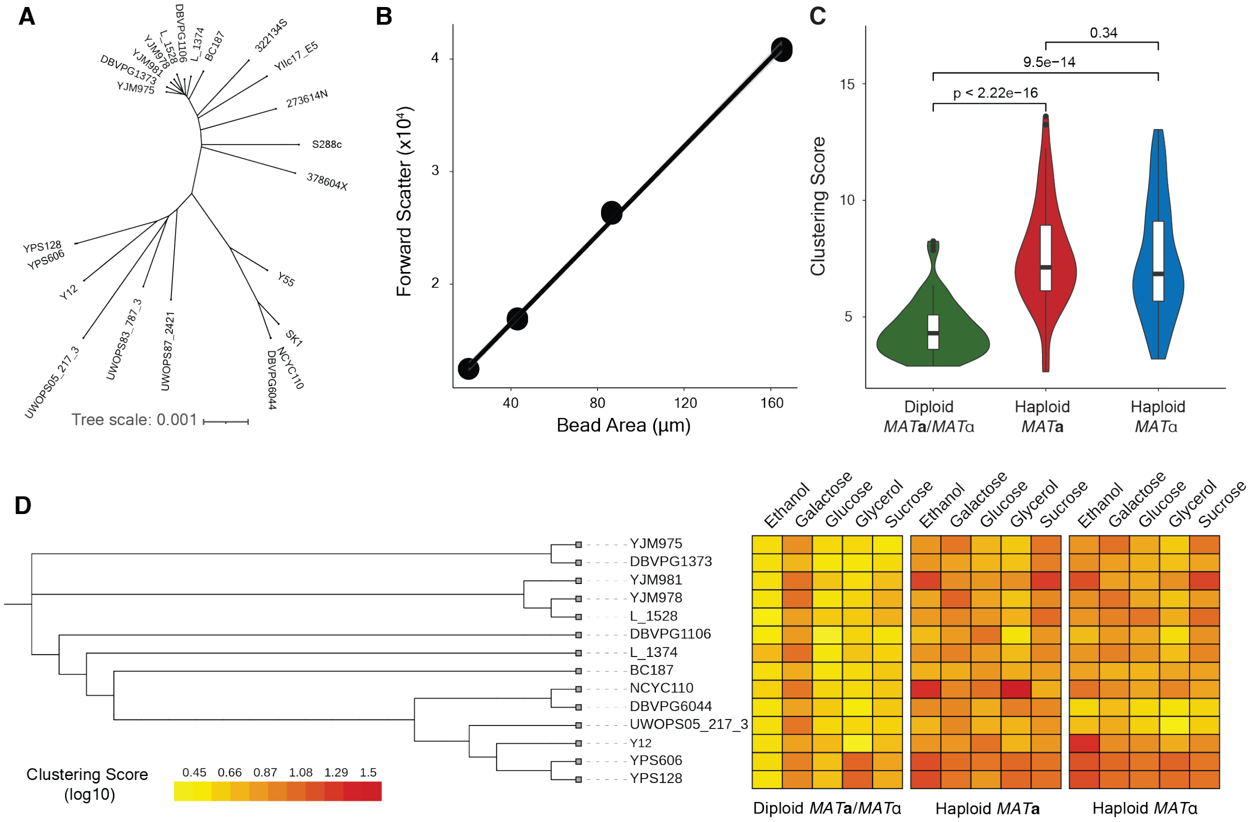


**Figure S1. Wild isolates phylogeny and clustering score. Related to Figure 1**

*A) Neighbor Joining tree of the wild isolates used in this study. Trees built based on SNP differences, using data from Liti et al. 2009[13].*

*B) correlation between bead cross-sectional area and mean forward scatter measurements. FSC for each bead size was measured in 3 independent measurements.*

*C) Clustering score in YPD of all 23 strains. The indicated p-values are the results of Wilcoxon test.*

*D) Clustering score in different carbon sources arranged by phylogeny. Data restricted to the 13 strains that are not mosaics from multiple lineages.*

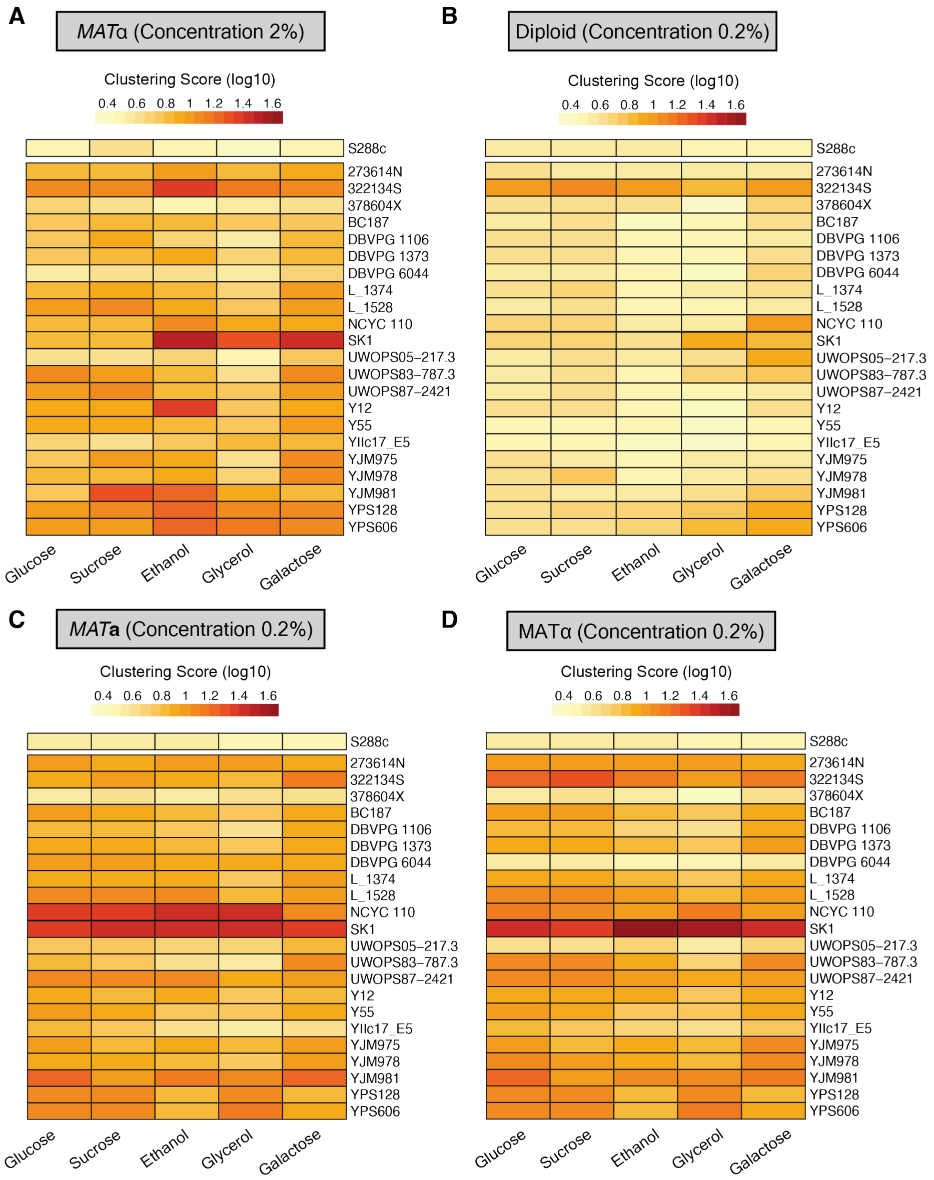


**Figure S2. Wild isolates clustering score at different sugar concentrations. Related to Figure 2**

A) *Clustering score for haploid MAT*$\alpha$ *isolates. Environments are hierarchically grouped. Strains were grown in YEP complemented with 5 different carbon source (concentration of 2% (w/v)). The clustering scores are the average of 3 measurements from independent experiments.*

B) *Same as A but for diploid strains in sugar concentration of 0.2%.*

C) *Same as A but for MAT**a** strains in sugar concentration of 0.2%.*

D) *Same as A but for MAT*$\alpha$ *strains in sugar concentration of 0.2%.*

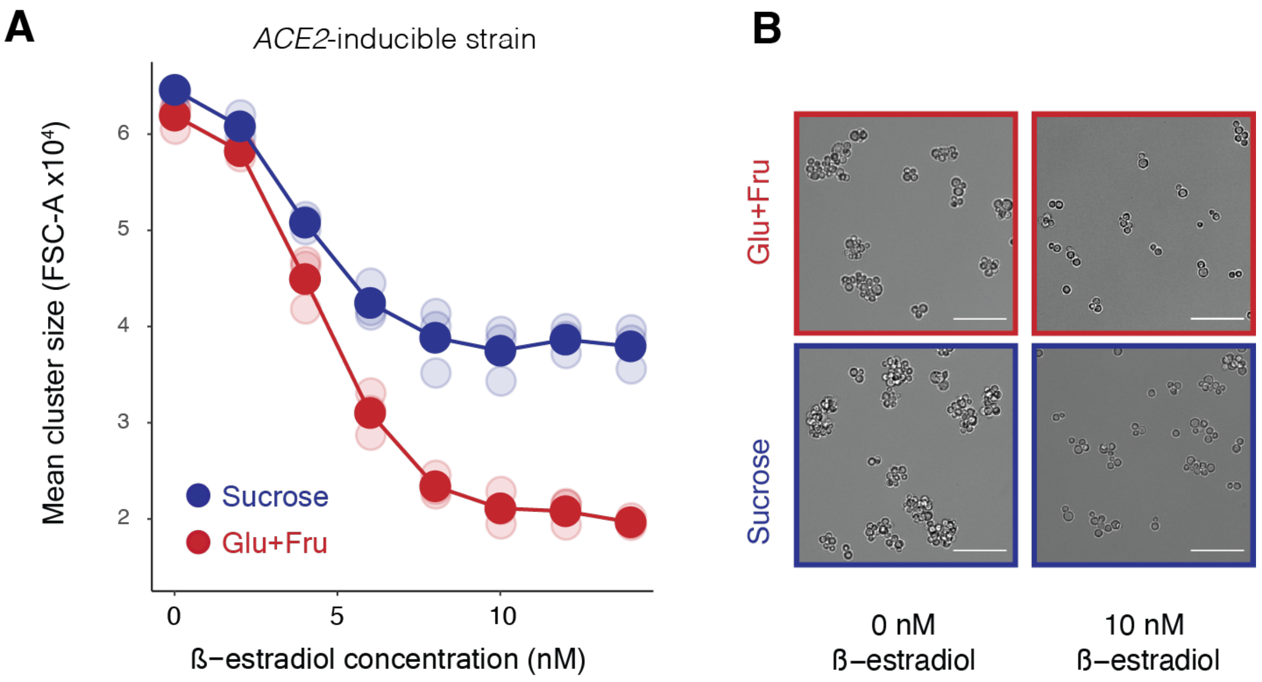


**Figure S3. Cluster size regulation using β-estradiol. Related to Figure 3**

*A) Mean forward scatter of the engineered, ACE2- inducible strain in glucose + fructose or sucrose at increasing concentration of β-estradiol. Error bars, too small to be seen, represent the standard error of the mean for 3 biological replicates.*

*B) DIC images of the ACE2-inducible strain when uninduced or induced with 10 nM of β-estradiol. Cells were grown in YNB with 10 mM sucrose or YNB with 10 mM glucose and 10 mM fructose. Scale bar represents 50 μm.*

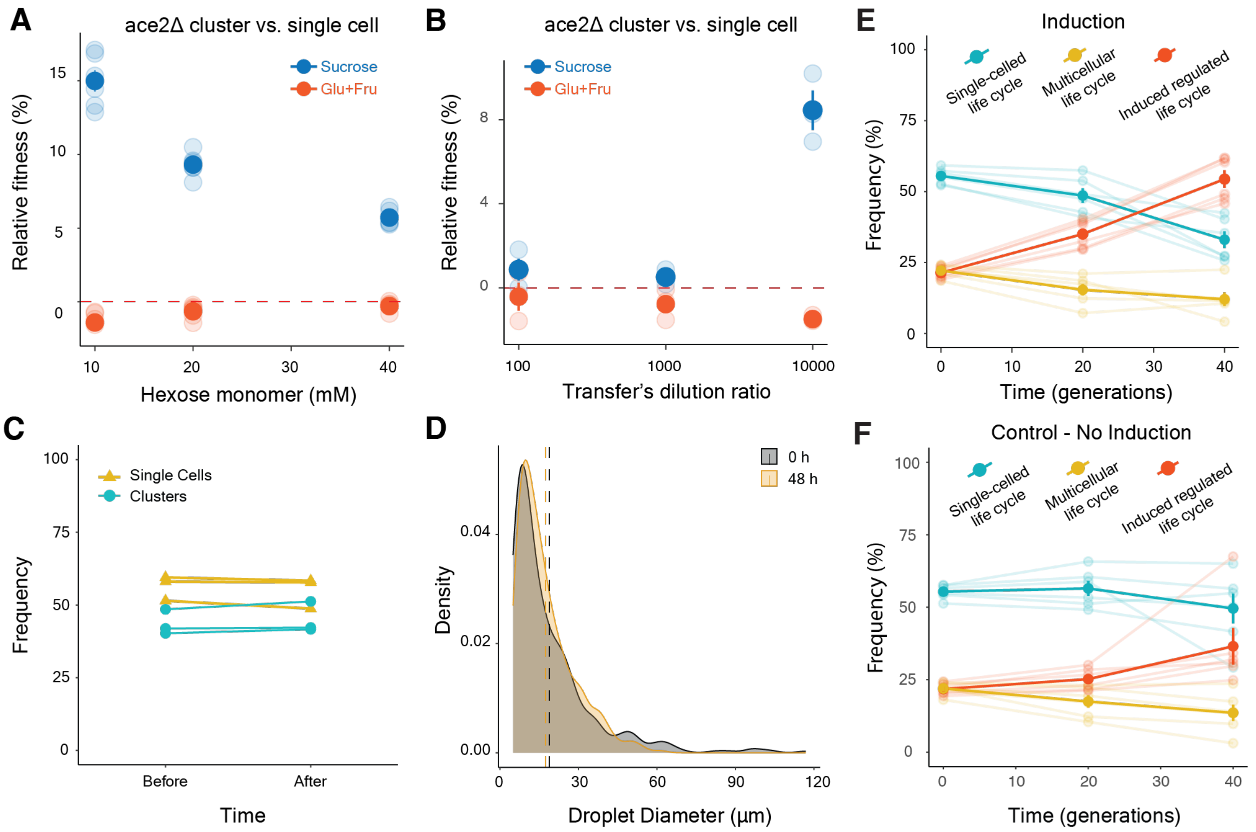


**Figure S4. Engineered life cycles can be competed in sucrose and glucose emulsions. Related to Figure 4**

*A) Competition experiment between an ace2Δ cluster and a single cell in different concentrations of sucrose or glucose + fructose. These results confirm work by Koschwanez et al. (2013) and quantify the fitness advantage of clusters in sucrose. Error bars represent the standard error of the mean for 3 biological replicates.*

*B) Similar to A but competitions were done in 10 mM sucrose and 10 mM Glucose + 10 mM Fructose with different dilution factors between transfers (100 stands for a 1:100 dilution). A large dilution factor is needed to observe fitness differences between single cells and clusters. Error bars represent the standard error of the mean for 3 biological replicates.*

*C) Measurements of ratio of single cells to clusters before introducing them into an emulsion and after immediately breaking the emulsion, with no intervening growth, to confirm that the emulsion protocol does not create a bias in favor of clusters or single cells.*

*D) Size distribution of emulsion droplets. The emulsion was imaged and droplet diameters were measured using Fiji.*

*E) Frequency of the three life cycles (single-celled, multicellular and regulated induced) overtime when all were grown together in a regime alternating between bulk sucrose and glucose emulsions. The bold lines represent the means of replicates, while the light lines represent the data for individual replicates. Error bars represent the standard error of the mean for the 6 biological replicates.*

*F) Same as E, but a control experiment where the strain designed as a regulated life cycle is not induced and therefore always grows as cluster.*

| Wild Isolate | Number of Counted Objects | Number Of Single cells | Number of Clusters | Percentage of Single Cells |
|---|---|---|---|---|
| 273614N | 157 | 31 | 126 | 19.7 |
| BC187 | 204 | 22 | 182 | 10.8 |
| L_1374 | 276 | 25 | 251 | 9.1 |
| NCYC 110 | 83 | 8 | 75 | 9.6 |
| S288c | 161 | 157 | 4 | 97.5 |
| SK1 | 114 | 11 | 103 | 9.6 |
| Y12 | 301 | 110 | 191 | 36.5 |
| YIIc17_E5 | 187 | 100 | 87 | 53.5 |
| YJM978 | 150 | 26 | 124 | 17.3 |
| YPS606 | 162 | 79 | 83 | 48.8 |

**Table S1: Percentage of single cells in wild isolates populations (MATa). Related to Figure 1**

*We randomly selected 10 wild isolates haploid MAT**a**, imaged them under the microscope and counted objects by classifying them as single cells or clusters.*

| Strain Name | Purpose | Genotype |
|---|---|---|
| yJB064 | Cluster forming strain | *can1-100 HIS3Δ::prACT1-yCerulean-tADH1-His3MX6 BUD4-S288C ACE2Δ::bleMX* |
| yJB077 | Reference strain for competition experiment | *can1-100 HIS3Δ::prACT1-ymCherry-tADH1-His3MX6 BUD4-S288C* |
| yJB128 | Single-celled strain with β-estradiol construct. | *can1-100 his3-11,15::PACT1(-1-520)-LexA-ER-haB42-TCYC1 BUD4-S288C HOΔ::prACT1-yCerulean-ADH1tr-KanMX* |

| | | |
|---|---|---|
| yJB130 | Inducible strain with β-estradiol construct. | *can1-100 his3-11,15::PACT1(-1-520)-LexA-ER-haB42-TCYC1 BUD4-S288C prACE2Δ:: PTEF-HygMX-TTEF-insul-(lexA-box)1-PminCYC1 HOΔ::prACT1-ymCherry-ADH1tr-KanMX* |
| yJB136 | Single-celled strain with β-estradiol construct. | *can1-100 his3-11,15::PACT1(-1-520)-LexA-ER-haB42-TCYC1 BUD4-S288C HOΔ::prACT1-ymCitrine-ADH1tr-KanMX* |
| yJB138 | Cluster forming strain with β-estradiol construct. | *can1-100 his3-11,15::PACT1(-1-520)-LexA-ER-haB42-TCYC1 BUD4-S288C HOΔ::prACT1-ymCitrine-ADH1tr-KanMX ACE2Δ::HphMX* |

**Table S2. Laboratory yeast strains used in this study. Related to Figure 3 and Figure 4**

*All strains were derived from yJK089, a derivative of the W303 strain background.*
